# Variability in codon usage in Coronaviruses is mainly driven by mutational bias and selective constraints on CpG dinucleotide

**DOI:** 10.1101/2021.01.26.428296

**Authors:** J. Daron, I.G. Bravo

## Abstract

The Severe acute respiratory syndrome coronavirus 2 (SARS-CoV-2) is the third virus within the *Orthocoronavirinae* causing an emergent infectious disease in humans, the ongoing coronavirus disease 2019 pandemic (COVID-19). Due to the high zoonotic potential of these viruses, it is critical to unravel their evolutionary history of host species shift, adaptation and emergence. Only such knowledge can guide virus discovery, surveillance and research efforts to identify viruses posing a pandemic risk in humans. We present a comprehensive analysis of the composition and codon usage bias of the 82 *Orthocoronavirinae* members, infecting 47 different avian and mammalian hosts. Our results clearly establish that synonymous codon usage varies widely among viruses and is only weakly dependent on the type of host they infect. Instead, we identify mutational bias towards AT-enrichment and selection against CpG dinucleotides as the main factors responsible of the codon usage bias variation. Further insight on the mutational equilibrium within *Orthocoronavirinae* revealed that most coronavirus genomes are close to their neutral equilibrium, the exception is the three recently-infecting human coronaviruses, which lie further away from the mutational equilibrium than their endemic human coronavirus counterparts. Finally, our results suggest that while replicating in humans SARS-CoV-2 is slowly becoming AT-richer, likely until attaining a new mutational equilibrium.

## Introduction

The *Severe acute respiratory syndrome coronavirus 2* (SARS-CoV-2) is the cause of respiratory disease COVID-19, occasionally leading to acute respiratory distress syndrome and eventually death (Zhu et al. 2020). With no antiviral drugs nor vaccines, and with the presence of asymptomatic carriers, the COVID-19 outbreak turned into a public health emergency of international concern. Before the 2019 zoonotic spillover, SARS-CoV-2 had circulated unnoticed for decades in bats as well as in other intermediate hosts, notably the Sunda pangolin *Manis javanica* (Boni et al. 2020). Cross-species transmissions are common among coronaviruses (CoVs), as all human infecting-CoVs (hCoVs) arose through zoonotic events, mostly from bats or rodents, with occasionally domestic animals playing the role of intermediate hosts (Cui et al. 2019). Consequently, given the high zoonotic potential of the *Orthocoronavirinae* (Olival et al. 2017; Dhama et al. 2020) and the vast repertoire of mammalian and avian hosts they infect, there is a need to evaluate the potential zoonotic risk for each individual CoV. This knowledge will guide virus discovery, surveillance and research to identify for each virus the differential risk to efficiently infect humans and cause a new pandemic.

Due to their highly compact genomes, none of the viruses infecting vertebrates encodes for any tRNA nor for any ribosomal protein, and CoVs are no exception. As a result, the translation of viral proteins relies exclusively on the host tRNA repertoire and translational machinery (Albers and Czech 2016). In order to efficiently support the production of viral proteins, it has been proposed that viruses should use the set of synonymous codons found overrepresented in their hosts, as the result of an adaptation to their host cellular environment (Franzo et al. 2017; Simón et al. 2017; Rahman et al. 2018; Tian et al. 2018). Such hypothesis is based on the fascinating discovery that codon usage is subject to natural selection (Ikemura 1985a; Kanaya et al. 2001; Drummond and Wilke 2008; Hershberg and Petrov 2008; dos Reis and Wernisch 2009). This adaptive hypothesis called translational selection proposes that the non-random usage of synonymous codons and the abundance of tRNAs have co-evolved to optimize translation efficiency. Experimental analyses have been successful to characterize how synonymous substitutions influence the cellular fitness of an organism, by acting on a broad range of cellular processes, including changes on transcription (Zhou et al. 2016), translation initiation (Kudla et al. 2009; Goodman et al. 2013), translation elongation (Sørensen et al. 1989), translation accuracy (Akashi 1994; Drummond and Wilke 2008), RNA stability (Presnyak et al. 2015), and splicing (Pagani et al. 2005).

Numerous works conducted essentially in certain phages and their bacterial hosts have reported a strong match between the codon usage bias (CUB) of viral genes with respect to their hosts (Lucks et al. 2008; Bahir et al. 2009; Wong et al. 2010). Similarly, the CUB of human papillomaviruses has been associated to differential clinical presentations of the infections (Félez-Sánchez et al. 2015), with a stronger virus-host match in CUB for human papillomaviruses causing productive lesions than those causing asymptomatic infections. However, such a trend has not been observed for SARS-CoV-2: most of the highly frequent codons in SARS-CoV-2 are A- and U-ending codons (Gu et al. 2020; Tort et al. 2020) overall showing a poor match with the average human CUB (Gong et al. 2020). Such mismatch has raised important questions on the nature of the CUB in CoVs, the underlying mechanisms involved, and the impact on virus longer-term evolution. Further, investigating the CUB match between viral and cellular genes constitutes a challenge to support the use of synonymous mutations as biotechnological tools to develop live attenuated vaccines (Lauring et al. 2010).

Besides been driven by constraints on tRNA abundance, CUB in viruses is also shaped by dinucleotide abundance. It has been long established that many RNA viruses infecting mammals have evolved CpG and UpA deficiency (Yap et al. 2003; Greenbaum et al. 2008; Atkinson et al. 2014; Takata et al. 2017). In the mammalian RNA Echovirus 7, it has been formally demonstrated that the severe attenuation of the virus replication is attributed to the increase of CpG and UpA dinucleotides frequencies rather to the use of rare codons (Tulloch et al. 2014). Viruses with high CpG/UpA frequencies may be more likely to be recognized by pathogen innate immune sensors, preventing them from replication initiation (Kumagai et al. 2008). Functionally, the inhibition of viral replication and the degradation of the viral genome have been attributed to the zinc finger antiviral protein (ZAP), a powerful restriction factor that specifically binds to CpG and UpA motifs (Odon et al. 2019). It is hence critical when investigating CUB to disentangle confounding effects of CUB and dinucleotide abundance.

Finally, non-adaptive evolutionary forces can also affect the viral genome composition. Because fitness differences associated to individual codons are very small, it requires very large population sizes (*e*.*g*. highly productive and highly prevalent viral infections) for natural selection to act and lead to a significant impact on the global genomic CUB. Such trend is verified in large organisms with small population sizes, such as most mammals, where natural selection on CUB is weak (Duret 2002; Chamary et al. 2006) and CUB is instead primarily shaped by mutational biases (Lynch 2010) and GC-biased gene conversion (Duret and Galtier 2009). Mutations are the fundamental substrate for genotypic diversity, leading to phenotypic diversity upon which natural selection can act. Point mutations are stochastic processes but they concur with certain deterministic and directional biases: in bacteria (Hershberg and Petrov 2010) as well as in eukaryotes (Petrov and Hartl 1999; Haddrill and Charlesworth 2008; Denver et al. 2009; Ossowski et al. 2010), previous studies suggest that mutations occurring during genome replication are universally biased towards AT. Similarly, for SARS-CoV-2, analysis of mutational profiles indicates a strong mutation bias towards U (Rice et al. 2020; Simmonds 2020), however the impact of such bias on CUB variation has not been characterized yet at the scale of the whole *Orthocoronavirinae*.

Given the impact of the match between virus and host CUB in viral gene expression, the key challenge is to determine whether a specific genome composition or CUB could facilitate (or hamper) the initial zoonotic spillover from animals towards humans and to eventually govern the risks of stable human-to-human transmission. Here we investigated the CUB variability in *Orthocoronavirinae*, with an emphasis on CoVs infecting humans. We aimed at determining whether CUB in CoVs is actively selected according to the codon preferences of their hosts or if it reflects the action of other evolutionary forces. We show that variation in CUB in *Orthocoronavirinae* mainly results from differences in GC3-content and in CpG and UpA abundance, independently of the host infected. Variation in GC3-content is primarily dictated by mutation biased towards AT, a trend universally observed in all *Coronavirinae* genera, regardless of the host. Finally, selection against CpG and UpA dinucleotides strongly impacts the CUB of CoVs. Altogether, we conclude that variation in CUB plays a little role, if any, on the probability of the establishment of a zoonotic spillover towards humans.

## Results

### Variation in codon usage bias among *Orthocoronavirinae*

To better understand the causes of CUB differences between CoVs, we investigated a total of 82 distinct CoVs. Our sample spans viral and host taxonomical diversity, covering the four viral genera within *Orthocoronavirinae*, and embracing a total of 47 different vertebrate host species within nine mammalian and five avian orders (Supplementary Table 1). First, we explored the variation in the 59 synonymous codons frequencies through a Principal Component Analysis (PCA) (Figure 1A and Supplementary Figure 1A). The PCA efficiently reduced information dimensionality as the first two components capture respectively 34.9% and 20.1% of the variance, and the first four dimensions contribute with above 70% of explanatory power. Interestingly, the 82 CoVs were distributed scattered along the two first PCA axes without any obvious stratification as a function neither of the viral taxonomy nor the host infected. This result contradicts our initial hypothesis of a host-specific CUB signature in CoVs and suggests that translational selection for convergence towards the host’s CUB is presumably weak.

**Figure 1:**
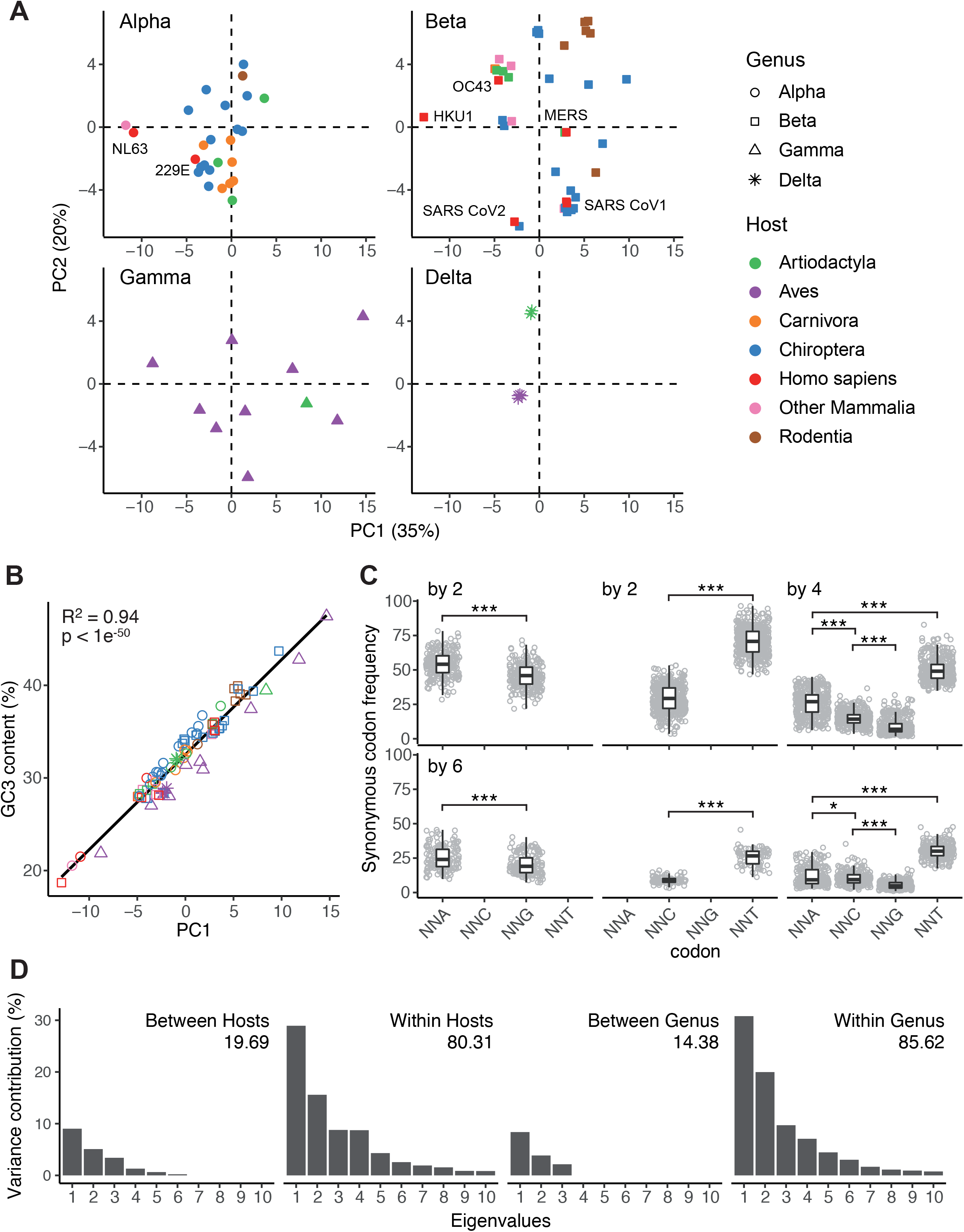
Variation in synonymous codon usage in *Coronavirinae*. A. Principal-component analysis of synonymous codon usage among CoVs. Each dot corresponds to an individual CoV, for which the frequencies of the 59 synonymous codon usage have been calculated. CoVs are stratified after the corresponding genera (Alpha-Delta), and host (colour code in the inset). The percentage of the variance explained by the first and second axes is given in parenthesis. B. Correlation between the projection on the first of principal components shown in (A) and the average GC-content at third codon position (GC3) for the corresponding viral genome. Symbol and colour code are the same as in (A). The results for a Pearson’s correlation test are shown in the inset. C. Synonymous codon usage frequency stratified by amino acids of multiplicity 2, 4, or 6 synonymous codons. Differences in median codon usage frequency were assessed by a paired Wilcoxon signed rank test (code for *p*-value: * <0.05; ** <0.01; *** <0.001). D. Results of an internal correspondence analysis for the contribution of the levels “viral taxonomy” (four levels, corresponding to viral genera) and “host taxonomy” (fourteen levels, corresponding to host order) to explain variability among the eigenvalues of viral genomes on the first ten principal components in (A). The values for expected explanatory power for a homogeneous distribution were and 3.72% (*p*-value < 9e^−4^) for viral taxonomy and 7.59% (*p*-value < 9e^−4^) for host taxonomy.

The PCA results contribute further with information about the spreading of the individual codons as well as their contribution to the total variance (Supplementary Figure 1B). The first PCA axis sharply splits the codons depending on their nucleotide in the third codon position (often referred to as GC3), at the exception of UUG-Leu codon, which stands alone among all other A- or T-ending codons. Strikingly, we found that variation in genomic GC3 content perfectly correlated with variation on the first PCA axis (R^2^ adjust=0.94, pvalue<1e^−50^, Figure 1B). Note that variation in GC3 did not show any correlation with any other of the main PCA axes (Supplementary Figure 2). In-depth analysis of the frequency patterns for the 18 synonymous codons families showed that A- or-U ending codons are systematically preferred over the G-or-C ending ones (Figure 1C). Such trend was especially true for amino acids with multiplicity-two, but also confirmed for amino acids with multiplicity four: U-ending codons were systematically the most used among the four codons, while A-ending codons were always preferred over the G-ending one. For amino acids with multiplicity six, the overall scheme corresponds to the combined patterns of a family of multiplicity four and a family of multiplicity two. Altogether, our observations show that variation in GC3 is the main explanatory factor for variation CUB between CoVs, with CoVs genomes being universally enriched in AT-ending over GC-ending synonymous codons.

We aimed at quantifying further the proportion of the global variability in CUB that is explained by the host and by the virus taxonomic stratification. Through a correspondence analysis, we associated the 82 different viruses of our codon usage table into blocks corresponding to the 13 host taxonomic orders, allowing us to split the total variability into between-category and within-category variability. The top ten eigenvalues of such decomposition are represented in Figure 1D showing that within-host differences in CUB explain four times more of the overall variability than differences between-hosts (respectively 80.3% *vs* 19.7%), the explanatory power of both levels being larger than the randomly expected one from a homogeneous distribution (7.59%, pvalue < 9e-4). A similar analysis was reproduced associating the viruses into blocks corresponding to the four viral genera within *Coronavirinae*. We observed that within-genus differences in CUB explain over five times more of the overall variability than between-genera differences (respectively 85.6% *vs* 14.4%), %), the explanatory power of both levels being again larger than the randomly expected one from a homogeneous distribution (3.72%, pvalue < 9e^−4^). Together, our results are consistent with our initial PCA analysis, and suggest that both host and virus taxonomy variables are not the main factors that drive variation in CUB within *Orthocoronavirinae*.

### The mutational spectrum in *Orthocoronavirinae* is AU-biased

Since variation in GC3 content is the main individual driver of the variation in CUB between CoVs, we investigated next whether mutational biases are the underlying main force driving nucleotide content. Population genetics studies show that in order to explore mutational biases independently of the effect of natural selection, one should work at shallow, short-term evolutionary periods, where natural selection is less powerful to impact nucleotide polymorphism patterns (Hershberg and Petrov 2010). Consequently, we estimated the frequencies of individual transition and transversions by narrowing down our analysis to a subset of 18 different CoV metapopulations. From the public databases (see Methods), we downloaded a total of 12,102 different viral genome entries to construct these 18 different metapopulations (Supplementary Table 2). Prior to the identification of SNPs for each metapopulation, we carefully preprocessed each dataset by removing putative recombinant sequences (see Methods, Supplementary Figure 3A), as those recombinant sequences violate the assumptions for the phylogenetic inference methods used downstream. For each viral metapopulation we removed further the effect of population structure by phylogeny-driven selection of one single homogeneous population per viral species (Supplementary Figure 3B). Our final dataset was composed of a total of 10,373 viral genomes for the 18 metapopulations (median number of genomes per metapopulation 31; range 9-9,588), for which a total of 57,059 SNPs were called (median number of SNPs called per metapopulation 3,037; range 253-7,778; Supplementary Table 2). Despite the heterogeneous sampling size across our metapopulations, the number of SNPs called was highly comparable, the exceptions being SARS-CoV-1 sequences from humans and MERS sequences from humans and from camels, which exhibited a low number of sequence polymorphisms, albeit large enough to grant sound analyses (respectively 716, 621 and 253).

Under a maximum likelihood framework, we separately estimated for the synonymous and non-synonymous compartments the frequencies of transitions and transversions using the generalized time-reversible model for the phylogenetic reconstruction fitting best the genomic data. Such analysis allows us to estimate the frequencies of individual transitions and transversions after correcting for nucleotide composition in each compartment. Our results show that for the synonymous substitutions compartment and for all viral metapopulations studied, U<>C transitions were more frequent than A<>G transitions, and U->C transitions were more frequent than the reverse C->U (Figure 2A). These remarkable differences were of a larger magnitude for substitutions occurring within the synonymous compartment than within non-synonymous compartment, where the C->U changes were only slightly more common than the reverse substitution. Consistent with these findings, stratification of substitutions into GC-enriching (*i*.*e*. AU->GC) or AU-enriching (*i*.*e*. GC->AU) categories shows that for all metapopulations AU-enriching substitutions occur at much higher rate than GC-enriching ones in both synonymous and nonsynonymous compartments (respectively mean fold change=2.49, *p*-value<7.6e^−6^; and mean fold change=1.38, *p*-value<7.6e^−6^; paired Wilcoxon signed rank test; Figure 2B).

**Figure 2:**
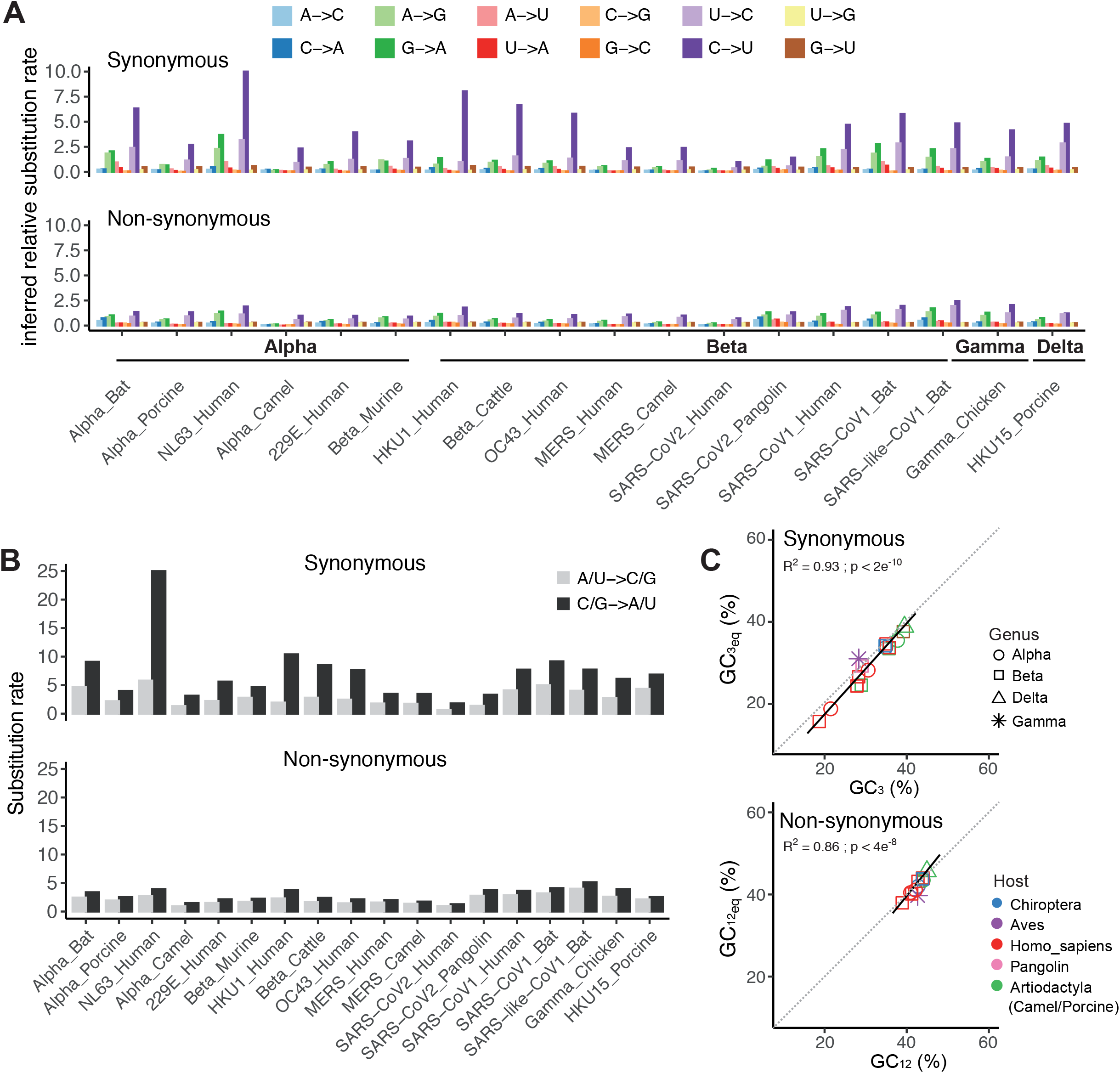
Mutational bias in CoVs. A. Relative substitution rates for all twelve transition and transversion events, estimated from the generalized time-reversible substitution matrix after maximum likelihood phylogenetic inference. For each virus the calculations were performed on the SNPs called separately on the synonymous or on the non-synonymous compartment of a metapopulation was composed with all available sequences, curated, filtered and aligned. B. Relative substitution rates for transition and transversion events resulting in GC enrichment (A/U->C/G) or in AT enrichment (C/G->A/T), inferred from data in (A) for the synonymous and non-synonymous compartments of the 18 viral metapopulations studied. C. (Upper panel) Plot of the observed GC composition in the third codon position (GC3) against the mutational equilibrium (GC_eq_) expected in the synonymous compartment. (Lower panel) Plot of the observed GC composition at first and second position of codons (GC_12_) against GC_eq_ estimated for the non-synonymous compartment. Symbols and color codes for the viral genomes are given in the inset. The continuous line corresponds to the regression line and the dotted gray line corresponds to the y=x plot. The values for the Pearson’s correlation are given for each panel.

Finally, we tested whether for each viral metapopulation the nucleotide content is at equilibrium in each compartment, by contrasting the observed GC content against the expected GC one if the metapopulation lay at its inferred mutational equilibrium (GCeq) (Figure 2C). We state first the very good correlation between observed and expected GC composition in both synonymous and non-synonymous SNPs (respectively R^2^ adjusted = 0.93 and 0.86; p-values < 2e^−10^ and 4e^−8^). Notably, CoVs infecting the same host displayed important differences in GC3% in the synonymous compartment, in the case of hCoVs ranging from 18.7% (HKU1) to 30.6% (229E). Such variation indicates again that the host is not the main factor governing the strength of the mutational bias. Altogether our results suggest that the mutational spectrum in CoVs is biased towards AU-enrichment and that the nucleotide content of the viral genomes is primarily determined by mutational biases.

### Recent human-infecting coronavirus display greater mutational disequilibrium than endemic hCoVs

Humans are the host to endemic CoVs, responsible for common respiratory diseases, but it has been proposed that these endemic hCoVs have an ancient zoonotic origin. Molecular dating studies have tried to evaluate the timing for host shift events for endemic hCoVs, estimating an old emergence of several hundreds of years ago for NL63 and a more recent one in the 19^th^-20^th^ century for 229E, OC42 and HKU1 (Vijgen et al. 2005; Pfefferle et al. 2009; Huynh et al. 2012; Al-Khannaq et al. 2016). This epidemiological stratification among hCoVs, differentiating ancient zoonosis that became endemic in humans and recently zoonotic hCoVs provided us with the unique opportunity to characterize the impact that spillover and subsequent establishment in the new host has on the mutational equilibrium of the virus genomes. Previous works have demonstrated that a high-resolution estimation of the mutational equilibrium can be gained from the simple property of the folded site frequency spectrum (SFS) of AT-to-GC mutations, in which SNPs are not polarized (Glémin et al. 2015). The theory proposes that for populations at demographic equilibrium the GC-content converges towards the mutational equilibrium, yielding a perfect U-shape in the AT-GC allele frequency distribution, resulting in a symmetrically folded SFS. Deviations from the U-shape distribution would thus reflect that the genomic GC-content is not at its mutational equilibrium, a negative (or positive) skewness indicating respectively higher (or lower) GC-content than expected under the mutational equilibrium. We have applied this framework, computed the folded SFS for the seven hCoVs lineages (Figure 3A) and quantified the deviation from the expected U-shape distribution by calculating the skewness of each folded SFS as a proxy for the departure of the GC-content from the expected mutational equilibrium (Figure 3B). Our results show that for all four endemic hCoVs the observed skewness was not different from the null expectation, while for the three recently emerged hCoVs (SARS-CoV-1, MERS-CoV and SARS-CoV-2) we observed a significant depart from the expected GC content at equilibrium. Both MERS-CoV and SARS-CoV-2 displayed a negative skewness, meaning that their genomes were slightly GC-richer than the anticipated mutational equilibrium, while SARS-CoV-1 displayed a positive skewness, reflecting an AT-richer genomic content than expected for the mutational equilibrium. Finally, benefiting from the wealth of sequence data generated on SARS-CoV-2, we investigated the dynamics of the GC-content over the spread of the COVID19 pandemic (Figure 3C). We observed that the GC-content of SARS-CoV-2 is very slowly albeit significantly decreasing with the progression of the pandemic (Figure 3C, R=-0.11, pvalue<1e^−49^). This result of a trend towards an overall AT-enrichment is consistent with the SFS observation of a viral genome far from compositional equilibrium and suggests that during the expansion of SARS-CoV-2 in humans, the virus population is experiencing a GC->AT mutational bias resulting in a slow decrease of the GC genomic content compared to the viral genomes retrieved from humans at the origin of the pandemic.

**Figure 3:**
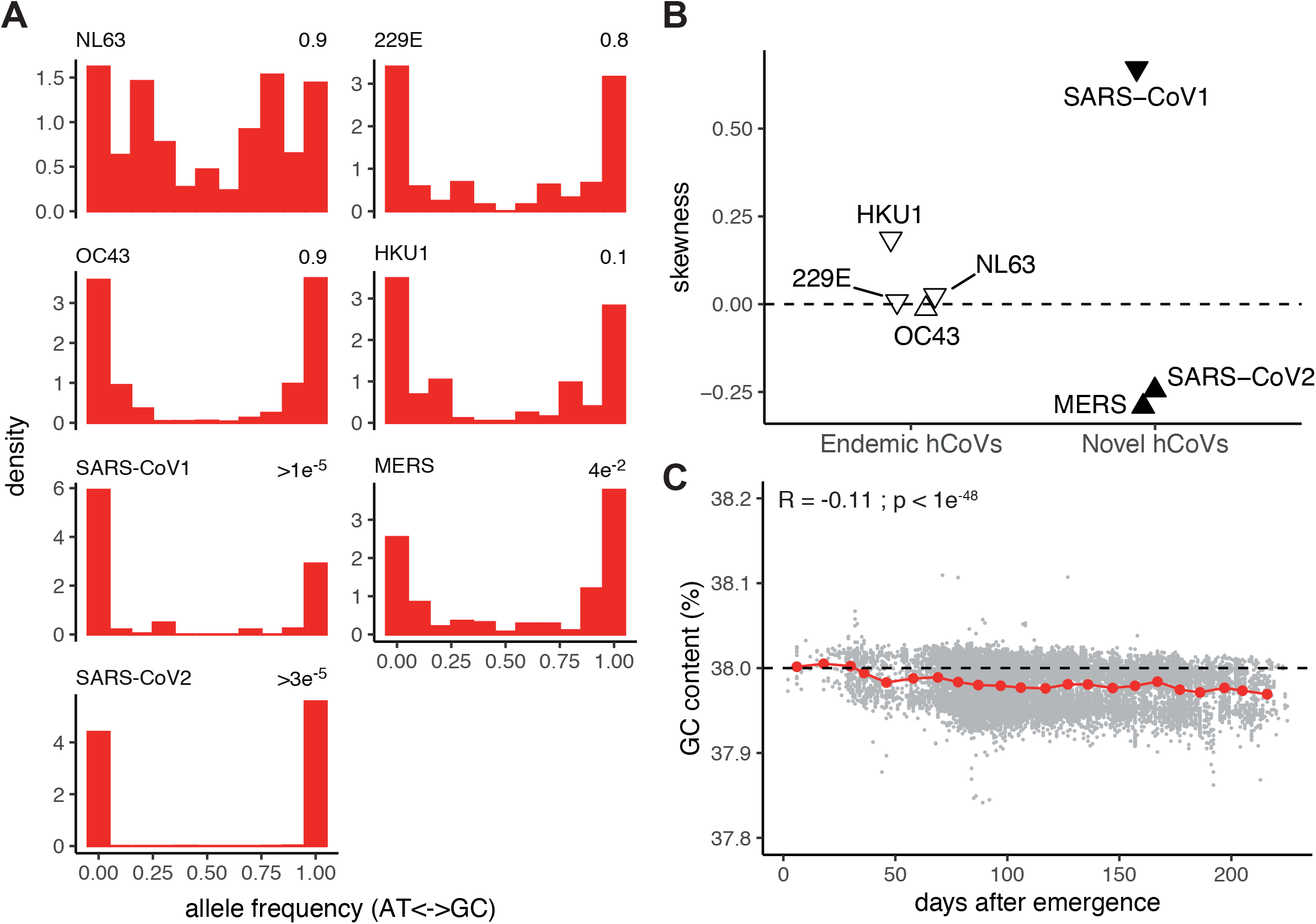
Mutational equilibrium among endemic and recently zoonotic human coronaviruses. A. Folded Site Frequency Spectrum (SFS) of G/C-to-A/T substitutions in the seven hitherto knew CoVs infecting humans. For each viral metapopulation the frequencies of SNPs involving a change in GC content called are plotted. For each viral metapopulation the probability that the skewness observed in the plot is different from the perfectly symmetrical distribution expected if the metapopulation was at its mutational equilibrium is given on the top-right. B. Comparison of folded SFS skewness presented in (A) between endemic and recently zoonotic CoVs infecting humans. The downward and upward triangle represented negative and positive skewness, respectively. The filled triangle represented metapopulations with a skewness significantly different from the expected one at equilibrium. C. Evolution of the SARS-CoV-2 GC-content in the coding genome over spread of the pandemic. The GC-content of each individual SARS-CoV-2 genome is represented by a gray dot. Red dots represent the average GC-content by time windows of ten days. Results of a Pearson correlation test are displayed in the inset, above for the complete data set (n=81963).

### CpG dinucleotides are selected against in *Orthocoronavirinae* genomes

Our initial characterization demonstrated that variation in GC3 content is the prevailing force shaping CUB among CoVs, explain one-third of the total variation in CUB. We have thus tried to identify other evolutionary forces further shaping CUB in CoVs. Previous works have identified that in several RNA viruses infecting humans certain dinucleotides are under-represented, notably CpG and UpA, and that this low dinucleotide frequency has a strong impact on codon-pair bias (Kunec and Osterrieder 2016). We investigated thus the ratio of the observed over the expected dinucleotide frequency in the CoVs coding sequences. Figure 4A shows that the observed/expected abundance of the CpG dinucleotide in all CoVs lineages is significantly lower than the null expectation based on the individual nucleotide frequency, while this is not the case for the UpA dinucleotide. To a lesser extent, we identify CpA and UpG to be marginally more frequent than expected, which can be linked to the strong decrease in CpG, as they correspond respectively to the transitions CpG->CpA and CpG->UpG. We further state that the observed/expected ratios for CpG and UpA correlated well with the second and third dimension of our PCA analysis, respectively (R^2^ adjusted = 0.67 and 0.44; Figure 4B and C). It is important to remind that these dinucleotide frequency values have been estimated for the complete viral coding sequence but are not limited to the codons themselves, *i*.*e*. we have also considered the presence of CpG and UpA dinucleotides in the codon boundary context, so that this impact is not simply related to higher frequency of CG-rich or AT-rich codons. This specific point will be addressed below. Overall, our results show that variation in CUB between CoVs is associated with variation in CpG and UpA dinucleotide content.

**Figure 4:**
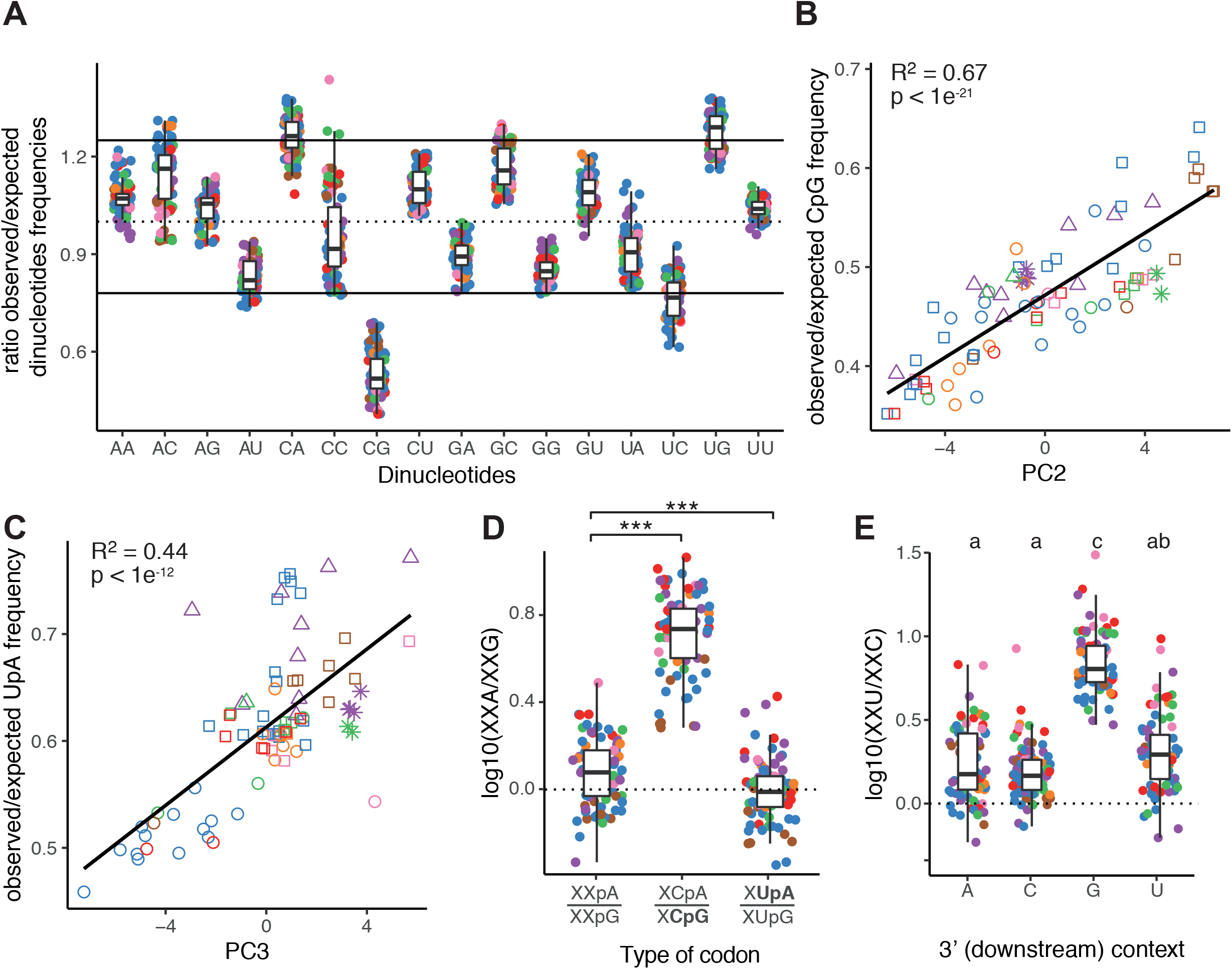
Dinucleotide Bias in *Coronavirinae*. A. Box (median, 1^st^ and 3^rd^ quartiles) and whiskers (95% CI) plot for the ratio of the observed over the expected frequency (dinucleotide relative abundance) for all 16 dinucleotides in the coding sequences of the 82 CoVs. Continuous lines indicate the thresholds for considering over/under representation of the corresponding dinucleotide over the expected values given the frequencies of the individual nucleotides. The threshold values at 0.78 and 1.25 have been phenomenologically determined following (Kunec and Osterrieder 2016). Colors correspond to the different host taxonomy orders. B. Linear regression between the projections of each individual viral genome on the second dimension of the synonymous codon usage Principal Component Analysis (Figure 1A) and the observed/expected frequency of the CpG dinucleotide. Shapes correspond to viral taxonomy genera and colours correspond to host taxonomy orders. C. Linear regression between the projections of each individual viral genome on the third dimension of the synonymous codon usage Principal Component Analysis and the observed/expected frequency of the UpA dinucleotide. Shapes correspond to viral taxonomy genera and colours correspond to host taxonomy orders. D. Box (median, 1^st^ and 3^rd^ quartiles) and whiskers (95% CI) plot for the ratio in synonymous codon frequency of A-over G-ending codons, calculated for codon families with multiplicity-two lacking CpG and UpA dinucleotides (Gln, Lys and Glu, indicated as “XXpA/XXpG”, constituting the reference set for comparison), amino acids encoded by CpG-ending codons (Ser4, Pro, Thr, Ala, indicated as “XCpA/XCpG”), and amino acids encoded by UpA-ending codons (Leu2, Leu4, Val, indicated as “XUpA/XUpG”). Colours correspond to the different host taxonomy orders. Differences within amino categories were assessed by a paired Wilcoxon signed rank test (Bonferroni correction, code for *p*-value: *** <0.001). E. Box (median, 1^st^ and 3^rd^ quartiles) and whiskers (95% CI) for the ratio in synonymous codon frequency in U-over C-ending codons, calculated for codon families with multiplicity four, and stratified by the nature of the 3’ downstream nucleotide. Differences depending on 3’ base context were assessed by a paired Wilcoxon signed rank test (Bonferroni correction) and summarized among sets of groups statistically different one from another. Colors correspond to the different host taxonomy orders.

Next, in order to formally demonstrate the impact that CpG and UpA dinucleotide frequency have on CUB, we compared the synonymous codon frequencies among codon families either containing or lacking a CpG- or UpA-ending codon. Having previously established that mutational bias modulates frequencies among synonymous codons, we accounted for this confounding effect by calculating an expected ratio in synonymous codon frequencies for codon pairs in the form of XXA/XXG. To this end we defined as our reference set for comparison the three codon families with multiplicity-two lacking CpG and UpA dinucleotides and ending by either A or G (*i*.*e*. Gln -CAA/G-, Lys -AA/G- and Glu - GAA/G-, indicated as “XXpA/XXpG” in Figure 4D). For this reference set, we calculated an overall fold change XXA/XXG ratio of 1.26, consistent with the AT-enriching mutational bias described above. For amino acids encoded by CpG-ending codons (Ser4, Pro, Thr, Ala, indicated as “XCpA/XCpG” in Figure 4D), we observed a 5.33-fold change XCA/XCG ratio, significantly higher than the corresponding one for the reference amino acids set (within-genome paired Wilcoxon signed rank test, *p*-value<1e^−14^). Such difference indicates that regardless of the amino acid encoded, CpA-ending codons are systematically preferred over their CpG-ending synonymous counterparts, at higher proportion than expected under the pressure of A->G mutational bias alone. In the case of UpA-ending codons (Leu2, Leu4, Val), we observed an overall XXA/XXG ratio of 1.04, slightly but statistically significantly lower than the corresponding one for the reference amino acid set (within-genome paired Wilcoxon signed rank test, *p*-value <1e^−6^). We interpret thus that although the UpA-dinucleotide is not significantly depleted in CoV genomes, UpA-ending codons are less frequent than their synonymous UpG counterparts. Consequently, the genomic UpA dinucleotide-content is shaped by two antagonistic evolutionary forces: on the one hand a mutational bias promoting the overall excess of G->A transitions and on the other hand a selection for XUpA->XUpG transitions in synonymous codons. Indeed, this interpretation supports again the finding of the UpG dinucleotide as being borderline significantly more frequent than expected, as it also corresponds to the transition UpA->UpG.

Finally, we aimed at characterizing the impact that the pressure against CpG and UpA dinucleotides imposes on CUB, when these dinucleotides are located at the codon pair boundary, such as XXC-GXX or XXU-AXX. We thus computed the frequency ratio XXU/XXC for synonymous codons depending on the nature on the downstream first nucleotide: if a selection again CpG nucleotides existed, we would expect the XXU/XXC ratio to be higher when the downstream codon starts with G; similarly, if a selection against UpA dinucleotides existed, the XXU/XXC ratio should be lower when the downstream codon starts with A. Our results confirmed indeed the depletion for CpG, as we observe a significantly higher XXU/XXC ratio upstream a codon starting by a G compared to codon starting with any other base (Figure 4E), avoiding the creation of CpG at the overlap between two codons. Regarding the hypothesis of UpA depletion, the median XXU/XXC ratio upstream a codon starting by A was the lowest of all bases, albeit with a large variance and not different from the values for codons starting by C or U. This result mitigates our previous finding of a slight intra-codon avoidance of the UpA dinucleotide, but is concordant with the overall absence of significant avoidance of UpA within the coding sequence.

### Mutational bias and CpG/UpA depletion explain most of the variation in synonymous codon usage of Coronaviruses

Having identified GC->AT mutational bias as well as CpG and possibly UpA dinucleotide depletion as variables impacting CUB in *Coronavirinae*, we have aimed at further quantifying the differential contributions of each of these factors on the overall CUB variation. We therefore built a linear model quantifying the relative contribution of the different variables (GC3, CpG, UpA, host taxonomy at the level of order, and virus taxonomy at the level of genus) and their corresponding interactions that covary with the projections of the first four main dimensions of the PCA for CUB. The analysis of variance demonstrated that variation in GC3, CpG and UpA are respectively the best predictors of the first three dimensions of the PCA (Figure 5), respectively explaining 94%, 67% and 56% of the variation of each dimension. By contrast, the integration of the host and virus taxonomy stratification levels and their interactions with the compositional variables did not contribute with any further improvement of the model fit to the data. Lastly, the fourth PC dimension was only marginally explained by UpA content and by the UpA*viral taxonomy interaction, which explained 19.3% and 18.2% of the PC4 dimension, respectively. Given these results and considering the power of each PCA axis for explaining variation in CUB, we conclude that variation in GC3 explains around 34%, variation in CpG frequency explains around 14%, and variation in UpA frequency explains around 7% of the total variation of CUB in *Coronavirinae*. On the contrary the contribution of viral taxonomy and of host taxonomy diversity to explain variation in CUB is negligible.

**Figure 5:**
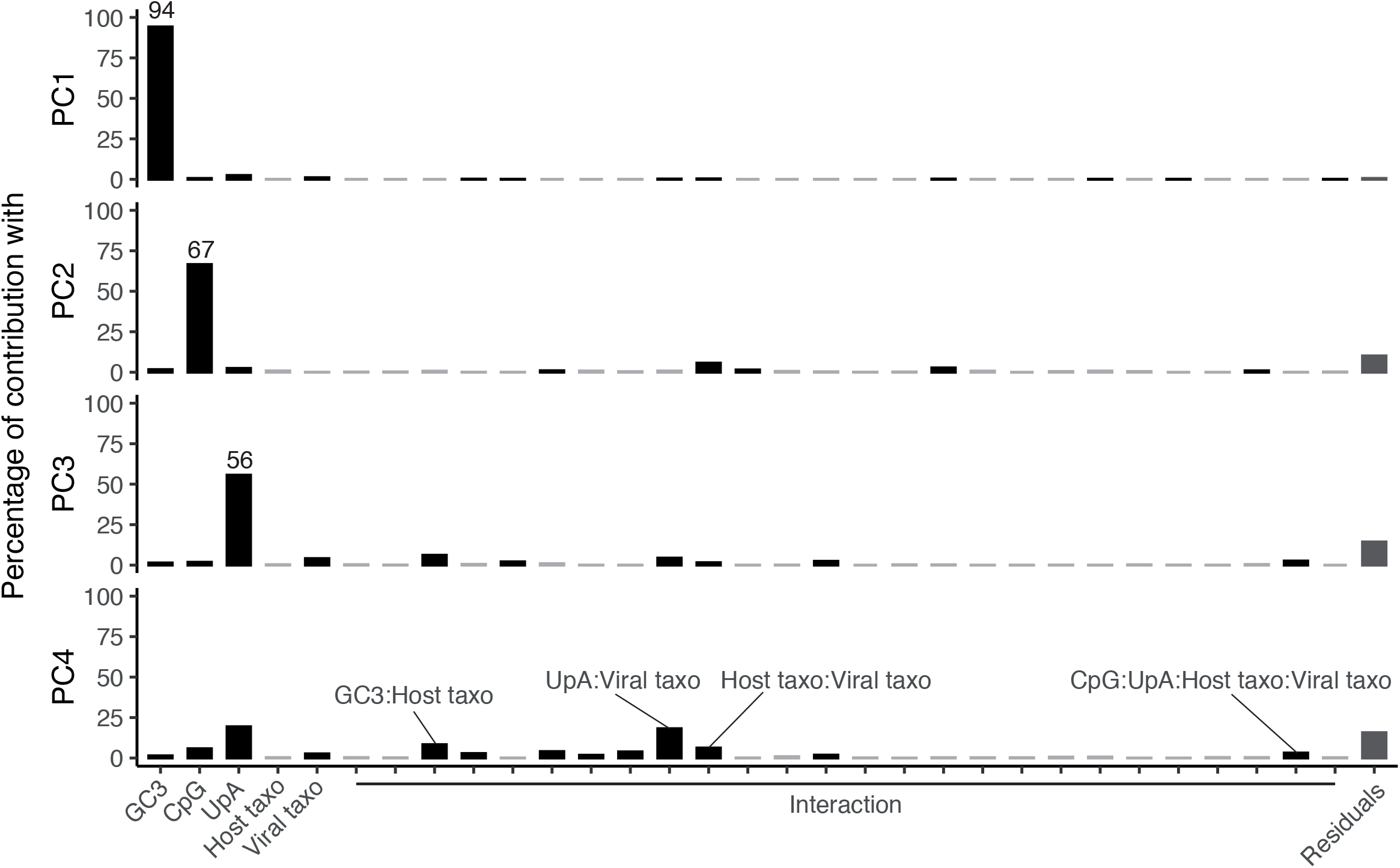
Linear model quantifying the relative contribution of the different variables to the projections of the first four main dimensions of the synonymous codon usage PCA (Figure 1A). Variables included in the linear model are: GC3-content, observed/expected frequency of the CpG dinucleotide, observed/expected frequency of the UpA dinucleotide, host taxonomy at the level of family, viral taxonomy at the level of genus, and all their pairwise interactions. Statistically significant contributions are shown as black bars, stating the percentage of the total explanatory contribution to the correspondent PCA dimension.

## Discussion

With the COVID19 pandemic, the unprecedented wealth of sequence data generated on human SARS-CoV-2 virus and the close relatives infecting pangolin and bats provides with a unique opportunity to investigate the variability and the biological role of synonymous codon usage among *Orthocoronavirinae*. Previous works in the field have been successful at inventorying CUB in SARS-CoV-2 (Dilucca et al. 2020; Gu et al. 2020; Tort et al. 2020), as well as at reporting the poor match in the observed CUB between SARS-CoV-2 and humans (Ji et al. 2020). Together those studies raised important questions on the nature of the CUB in CoVs, the underlying evolutionary mechanism involved, and their impact on the virus longer-term evolution. Here we explored those questions, by meticulously disentangling the effects that natural selection, dinucleotide avoidance and neutral forces have on the variability in CUB in CoVs.

### Mutational bias and CpG avoidance shape codon usage bias in CoVs

We report a strong heterogeneity in CUB among CoVs mainly driven by a GC->AT biased mutations. During analyses restricted to the SARS-CoV-2 genomes, previous studies had identified C->U deamination as the main mutational contributor to spontaneous mutation (Rice et al. 2020; Simmonds 2020). Our work conducted on the largest diversity of CoVs investigated so far is a novel compelling piece of evidence demonstrating that mutational bias toward AT is a universal feature among CoVs, regardless of the viral taxonomy and of the host infected. We further report that for all CoVs, C->U substitutions occur at a much higher frequency than G->A substitutions. Such asymmetric substitution pattern has been proposed to be driven by a host APOBEC3-like editing process, rather than occurring during virus replication and being associated to mutational biases of the coronavirus RNA-dependent RNA polymerase (Di Giorgio et al. 2020; Simmonds 2020). The *APOBEC3* locus in mammals consists of a series of tandem copies of different *APOBEC3* genes that have undergone a complex evolutionary history of duplications, deletions and fusions, further complicated at the transcriptome level with a large diversity of mRNA (Münk et al. 2012). The evolution of the *APOBEC3* locus is tightly linked to the viral infections via different arms race (Harris and Anderson 2016; Ito et al. 2020). The structure and synteny in the locus and the repertoire of APOBEC3 proteins encoded by one genome are thus extremely lineage-specific: see for instance (Hayward et al. 2018; Jebb et al. 2020) for Chiroptera or (Garcia and Emerman 2018; Yang et al. 2020) for Primates. It is thus important to note that the mutagenic role of host APOBEC3 enzymes acting onto viral genomes and exerting strong evolutionary pressures in a particular virus-host interaction may fuel viral adaptations, which facilitate viral transmission and colonisation of novel host species, as suggested for lentiviruses infecting Primates (Nakano et al. 2020).

In addition to the increased GC->AT substitutions, we remark that CoVs genomes are strongly depleted in the CpG dinucleotide, resulting in the avoidance of synonymous codons and of codon pairs containing this motif. Our results are consistent with previous works demonstrating that the experimental increase of CpG frequencies impaired virus fitness (Lauring et al. 2012; Tulloch et al. 2014; Moratorio et al. 2017). CpG depletion has been observed across CoVs infecting a whole range of mammals and birds, indicating that the attenuation mechanism might be fundamental to vertebrate eukaryotic antiviral defense, evolutionarily conserved and active over three hundreds of millions of years of evolution (Ibrahim et al. 2019). Surprisingly, we did not observe any statistically significant depletion of the UpA dinucleotide in CoV genomes (Figure 4A), while underrepresentation of UpA is a common trend among human RNA virus (Simmonds et al. 2013; Kunec and Osterrieder 2016). Analogous to the host immune response to the presence of CpG in the viral genomes, the increased presence of UpA dinucleotides has been formally identified as preventing virus replication (Fros et al. 2017), potentially through the cleavage of viral RNAs by RNase L (Cooper et al. 2015). Our investigation of the impact of UpA dinucleotide on the synonymous codon frequency displayed mitigated results: avoidance of UpA dinucleotide was observed when the UpA dinucleotide is located at the end of a codon (Figure 4D) but not when the UpA dinucleotide is located at the codon pair boundary (Figure 4E). Hence, we hypothesize that UpA sites in CoVs are at the center of the antagonist effects of mutation bias (promoting the formation of new UpA sites through GC->AT mutational bias) and selection (directly acting against UpA sites). However, testing such mutation-selection trade-off hypothesis goes beyond the power of our bioinformatics analyses and should be properly addressed by actually modifying the UpA content of CoV genomes, quantifying changes in viral fitness and following the evolutionary trajectories of the modified genomes by means of experimental evolution. Altogether, our work fits well with the mounting evidence supporting the biotechnological application of modifications in CpG and UpA dinucleotide frequencies in viral genomes for the production of efficient, safe and evolution-proof vaccines (Martínez et al. 2016; Stauft et al. 2019; Konopka-Anstadt et al. 2020).

### Lack of evidence for translational selection acting on CoVs

Mismatches between the mRNA characteristics and the translation machinery can have a strong impact on the quality and quantity of protein production. Selection acting on mutations leading to streamlined translation is known as translational selection. However, studies on the distribution of fitness effect show that the selective advantage associated to beneficial individual synonymous mutations is very low (Sanjuán et al. 2004; Peris et al. 2010; Jacquier et al. 2013; Fragata et al. 2018; Williams et al. 2020), implying that translational selection could only be efficient in the case of organisms with large effective population sizes. Indeed, strong phenotypic effects of codon usage on expression levels of single genes have been experimentally reported in some metazoan species with large population sizes (such drosophila or nematode) (Chamary et al. 2006; Plotkin and Kudla 2011). Similarly, studies in vertebrates have established that translational selection is much weaker in large-sized vertebrates compared to small-sized, because there are slow growing organism with a small effective population size (Galtier et al. 2018). Additionally, the physicochemical organization of vertebrate chromosomes may further hamper translational selection from playing a strong role in vertebrates: chromosomes in vertebrates are organized in long, consecutive regions enriched in AT or in GC nucleotides, known as isochores (Caspersson et al. 1968), so that the factor with the largest individual impact on CUB of vertebrate genes is its physical location along the chromosome (Holmquist 1989).

Just like cellular genes, viral genes rely completely on the host cell apparatus for translation. It has thus been proposed that selection could act to optimize the match between the virus CUB and tRNA repertoire and abundance in its host, leading to increased efficiency and accuracy of viral protein synthesis (Tian et al. 2018). In the case of viruses, the large population sizes that many of them generate during productive infections could allow translational selection to shape CUB. For bacteriophages, CUB has been shown to match codon preferences in the bacterial host (Lucks et al. 2008), which in their turn match the most abundant cognate tRNAs available in the cell (Rocha 2004). In viruses infecting vertebrates, mounting experimental evidence suggests that modification of viral codon usage leads to sharp changes in viral fitness, the necessary condition for natural selection to act upon viral CUB (Martínez et al. 2016). However, the results here presented show that host stratification explains only a minor fraction of the global variability of CUB in CoVs, suggesting that for these viruses the variability of synonymous codons is largely unrelated to the type of host infected. We conclude thus that, translational selection based on a differential tRNA abundance as a function of the host is not the main evolutionally driver of CUB in CoVs. Experimental results on ribosome profiling for *Murine coronavirus* genes have reported that despite the poor match in CUB between this virus and the mouse host, coronavirus mRNAs were translated with similar efficiency than the host mRNAs (Irigoyen et al. 2016). Thus, the high frequency of U- and A-ending synonymous codon in CoVs genomes, systematically departing from the CUB of their host (Ji et al. 2020), would not negatively impact the synthesis of viral proteins.

In metazoans the interplay between the main factors shaping CUB, mutational forces, GC-biased gene conversion and translational selection, have been finely analyzed and interpreted as a function of effective population size (Ratnakumar et al. 2010; Galtier et al. 2018). Notwithstanding, the same studies identify CUB trends, such as the systematic preference of pyrimidines over purines at the third codon position for which we still lack an interpretative framework (Galtier et al. 2018). In our analyses for CoVs, despite the strong explanatory power of variation in nucleotide composition and CpG avoidance to explain variation in CUB, over 40% of the overall variation in CUB still remains to be explained. Given the impact of effective population size on the efficiency of translational selection to shape CUB in metazoans, could translational selection contribute to explain the proportion of the variation in CUB in CoVs that has not been attributed to any variable? Here we report that the between-host variability of CUB in CoVs accounts for 19.7% of the total variability in CUB, a fraction two times larger than the random expectation. Although statistically significant, this contribution does not necessarily prove that differential adaptation to the translational machinery of the different hosts is shaping CUB in CoVs. Our analyses suggest instead that significant differences in GC3 and UpA occur between CoVs infecting different hosts (Supplementary Figure 7 and 8), so that between-hosts differences in CUB in CoVs could actually reflect again the impact of compositional variation at shaping CUB. A proper answer to the question of the actual role of translational selection in CoVs will require experimental work identifying differences in the natural history of the infection (*e*.*g*. target tissue, productivity, clinical presentation) of genetically close viruses upon host switch.

### Composition of CoVs genomes tends to reach their mutational equilibria

In order to investigate mutational bias in CoVs and to disentangle it from the effect of natural selection, we have analyzed mutations occurring during recent evolution in 18 viral metapopulations at different taxonomic host and viral levels. With this approach we have aimed at comparing mutational biases and equilibria, while minimizing the impact of natural selection. Notwithstanding, for all metapopulations the substitution rate in the synonymous compartment was systematically higher than in the corresponding non-synonymous compartment (Figure 2A-B). This result suggests that natural selection is still able to prune deleterious mutations at shallow levels in CoVs. We hypothesize that such an effect of natural selection may be linked to the viral life cycle allowing for within-host competition between putatively generated variants, and/or to the presence of bottlenecks or of differential variant success during within-host and during between-host transmission. This verbal argument can obviously be addressed only through observational data, quantifying the within-host viral diversity during the course of the infection, and its connection with the observed between-host viral diversity, or through experimental data using for instance mutation accumulation lines.

This metapopulation approach has allowed us also to address the extent to which each of the 18 CoVs may have reached a mutational composition equilibrium. Our results for all metapopulations show a very good correlation between the observed and the expected GC3 contents at both the synonymous and non-synonymous compartments, indicating that all CoVs were found at their GC-content equilibrium (Figure 2C). However, when considering the more accurate folded site-frequency spectrum at the population level, we observe that endemic hCoVs are closer to an equilibrium in the AT—GC allele distribution than zoonotic, recently acquired hCoVs (Figure 3A-B). This suggests that endemic hCoVs may have undergone a compositional drift towards a novel equilibrium under the novel mutational pressures in the human hos upon host switch. Indeed, at the short time scale that our analyses can explore for the SARS-CoV-2 epidemics in humans, we verify a small albeit significant trend towards GC reduction (Figure 3C). Future work will benefit from accessing a sufficient number of SARS-CoV-2 non-human strains collected from different hosts (bats and pangolins in first instance, given the current knowledge) to assess the population-level characteristics of the direction and intensity of mutational bias in endemic host species. Additionally, endemic hCoVs exhibit a large variation in observed synonymous GC3-content, ranging from 18.7% to 30.6%, which suggests that the strength of the mutational bias causing C-to-U change is not similar among all hCoVs. We hypothesize that such differences in mutational signature could be related to the differences in the host mutator APOBEC3 repertoires, which vary between hosts, but also between cell types within an organism, so that changes in the host tropism will have an impact on the actual mutational intensity and direction that the viral genomes experience.

## Conclusion

### CUB is a poor proxy to predict zoonotic infection in CoVs

SARS-CoV-2 is the third *Coronavirinae* zoonotic spillover, and the associated COVID19 poses an unprecedented burden on human public health. Understanding the drivers and facilitators of interspecies viral transmission in CoVs is a key public health and fundamental research priority. Nucleotide composition and CUB are distinctive characteristics of a viral, shaped by the global action of different pressures, including host adaptation (Jenkins and Holmes 2003; Wong et al. 2010; Cristina et al. 2015). In the case of SARS-CoV-2, the viral CUB is closer to that of humans than other hCoVs (Dilucca et al. 2020), which may have facilitated the establishment in humans following the zoonotic transmission event. Here we have shown that the main drivers of CUB (low GC3 content, strong CpG avoidance and slight UpA avoidance) are common to all CoVs, independently of viral taxonomy and of the hosts they infect. We interpret that the role of CUB at modifying the chances of a CoV zoonotic spillover to thrive in humans is negligible. Notwithstanding, the available data suggest that upon colonization of humans endemic hCoVs have experienced a compositional drift towards a novel compositional equilibrium in the new hosts, and that this could also be the case for SARS-CoV-2 if the current pandemics transforms into an endemic circulation in humans.

## Material and Methods

### Data collection and processing

To access the diversity of coronavirus species we downloaded sequences classified within the *Orthocoronaviridae* families from the Virus-Host database. Because the 212 virus entries present in the database are composed of a mixture of sequence classified at different taxonomy levels, *i*.*e*. species (HKU1, HKU4, HKU5,…) or strains (HKU4-1, HKU4-2, HKU4-3,…), we used CD-HIT (option –c 0.95 –n 5) to cluster together sequences sharing over 95% of identify and selected from each group one single representative strain.

For the intra-species diversity, we downloaded strains sequences from the Virus Pathogen Resource database (VIRP). Sequences unusually long or short (>130% or <70% of the median length of the reference sequence of a species) were filtered out. In addition, the complete genomic sequences of SARS-CoV-2 isolates were obtained from GISAID (available at https://www.gisaid.org/epiflu-applications/nexthcov-19-app/), accessed twice, on 2020/04/26 to collect the 9774 sequences for our metapopulation analysis and on 2020/08/18 to investigate the GC-content dynamics over the spread of the pandemic. Detailed accession ID for both datasets are provided in Supplementary Table 1 and Supplementary Table 2.

### Nucleotide Composition Analysis

We concatenated the coding sequences of each CoV genome in order to compute the total synonymous codon frequencies and GC-content. This analysis yielded a matrix of 82 (CoVs) x 59 (synonymous codons) which served as input for either our PCA analysis (FactoMineR) and correspondence analysis (ade4). In parallel, we calculated the dinucleotide observed/expected abundances in coding sequences by calculating the ratio rXY = fXY/fXfY, where fXY denotes the observed frequency of the dinucleotide XY and fXfY the product of the individual frequencies of the nucleotides X and Y in a sequence.

We used as significant lower and upper boundaries the thresholds of 0.78 and 1.25, corresponding to *p*<0.001 for sufficiently long (>20kb) random sequences (Burge et al. 1992).

### Sequence processing and SNP extraction

Prior to the SNP calling analysis, we identified and removed recombining sequence from each dataset using fastGear (Mostowy et al. 2017). Short recombination segments (<200 bp) identified by fastGear were not remove but those region were be mask in the downstream analysis. Next, in order to work in a framework where population structure will not interfere with our downstream analysis, for each species we generated a phylogenetic tree root with an outgroup and selected monophyletic group having the most of sequence for our downstream analysis. After those preprocessing sets, we finally performed using mummer (Delcher et al. 2002), a genome-to-genome alignment on each virus species, creating a pairwise alignment of each genome with a chosen reference, and performed a SNP calling analysis. Finally, ML phylogenetic trees and substitution matrix were inferred with RAXML (Stamatakis 2014), using the General Time Reversible (GTR) with an ascertainment bias correction model.

## Supplementary Figure and Table Legends

Supplementary Table 1: *Orthocoronaviridae* sequence data sets used for the study

Supplementary Table 2: Summary statistics of the 18 metapopulation of *Orthocoronaviridae*.

**Supplementary Figure 1:**
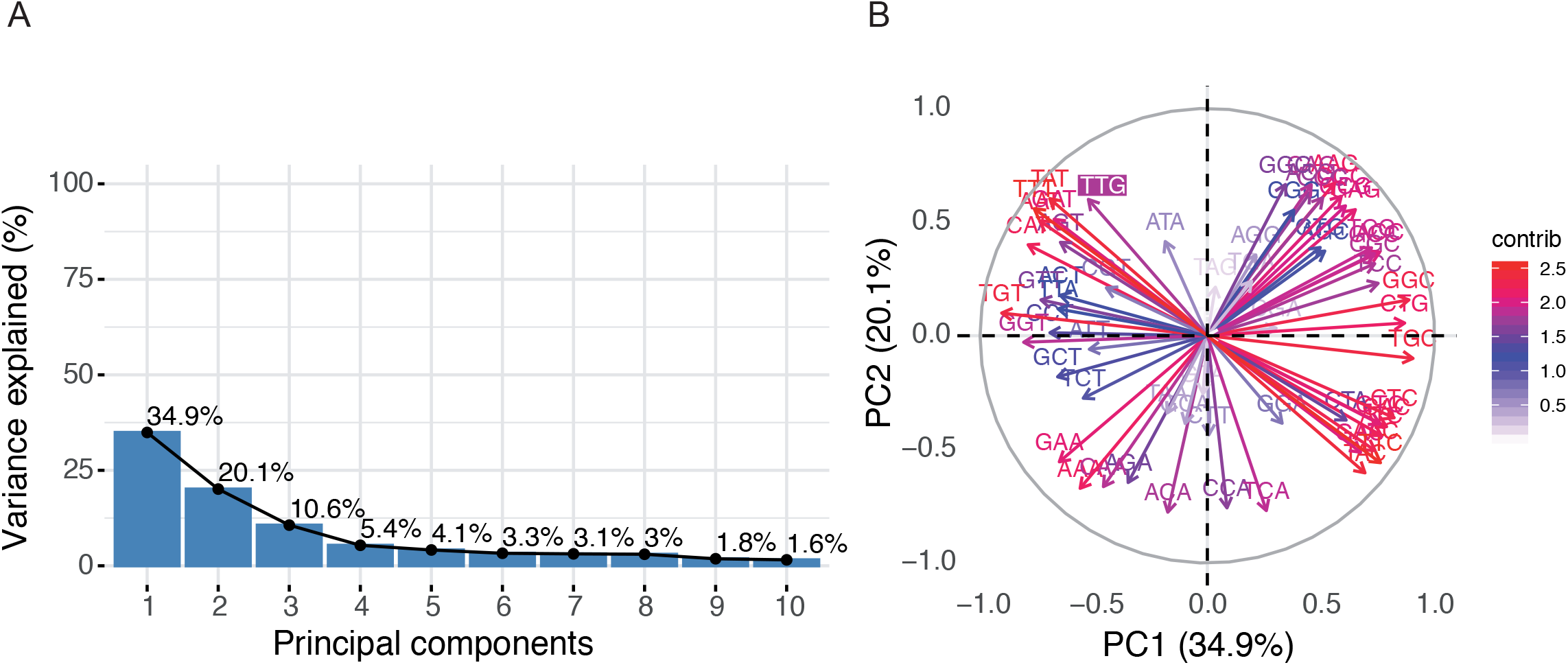
(A) Bar chart representing the percentage of variance explained by each principal component. (B) Projection of the individual codon on the first two principal components.

**Supplementary Figure 2:**
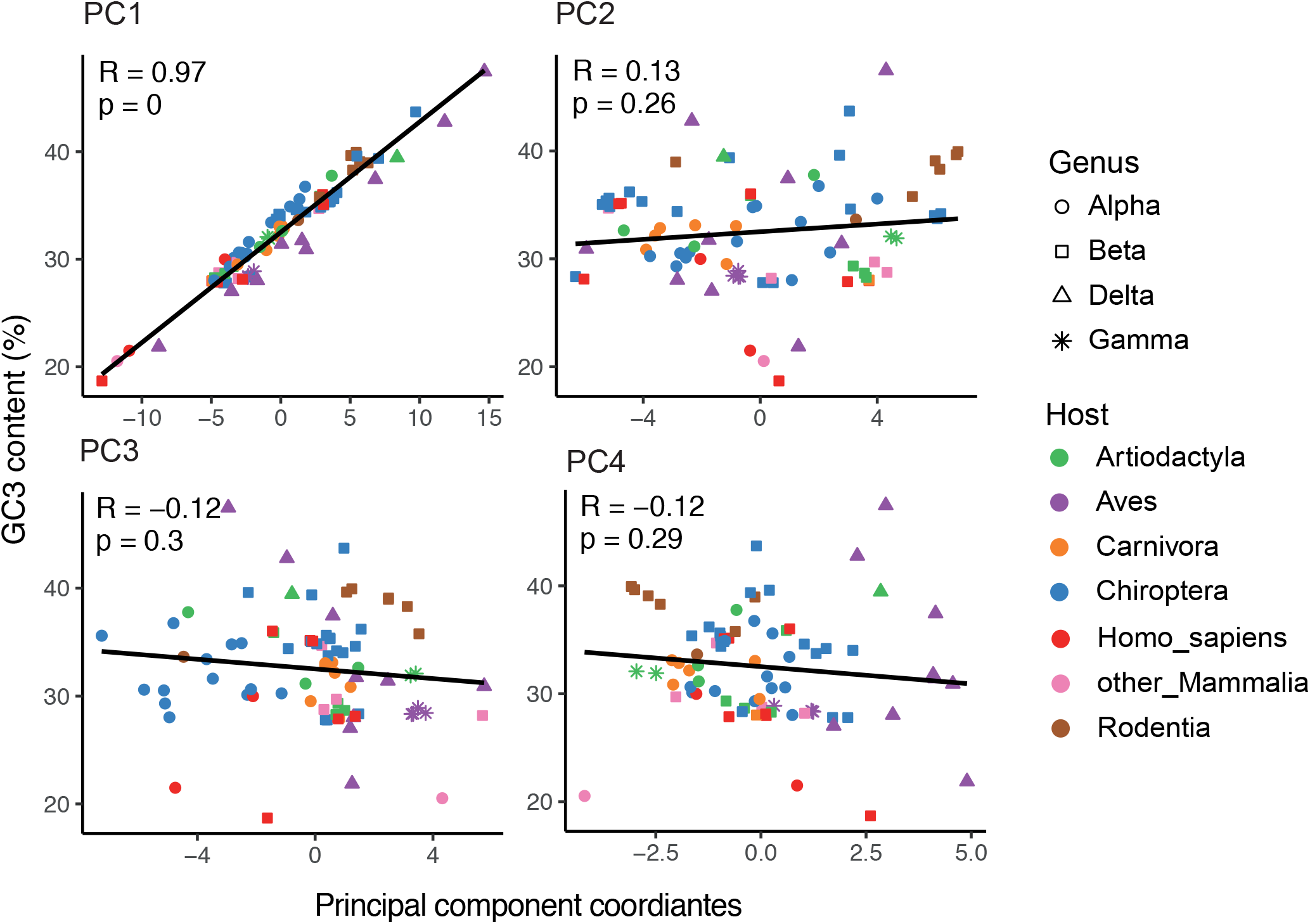
Regression between GC3 content (%) and coordinate of the first 4 principal components.

**Supplementary Figure 3:**
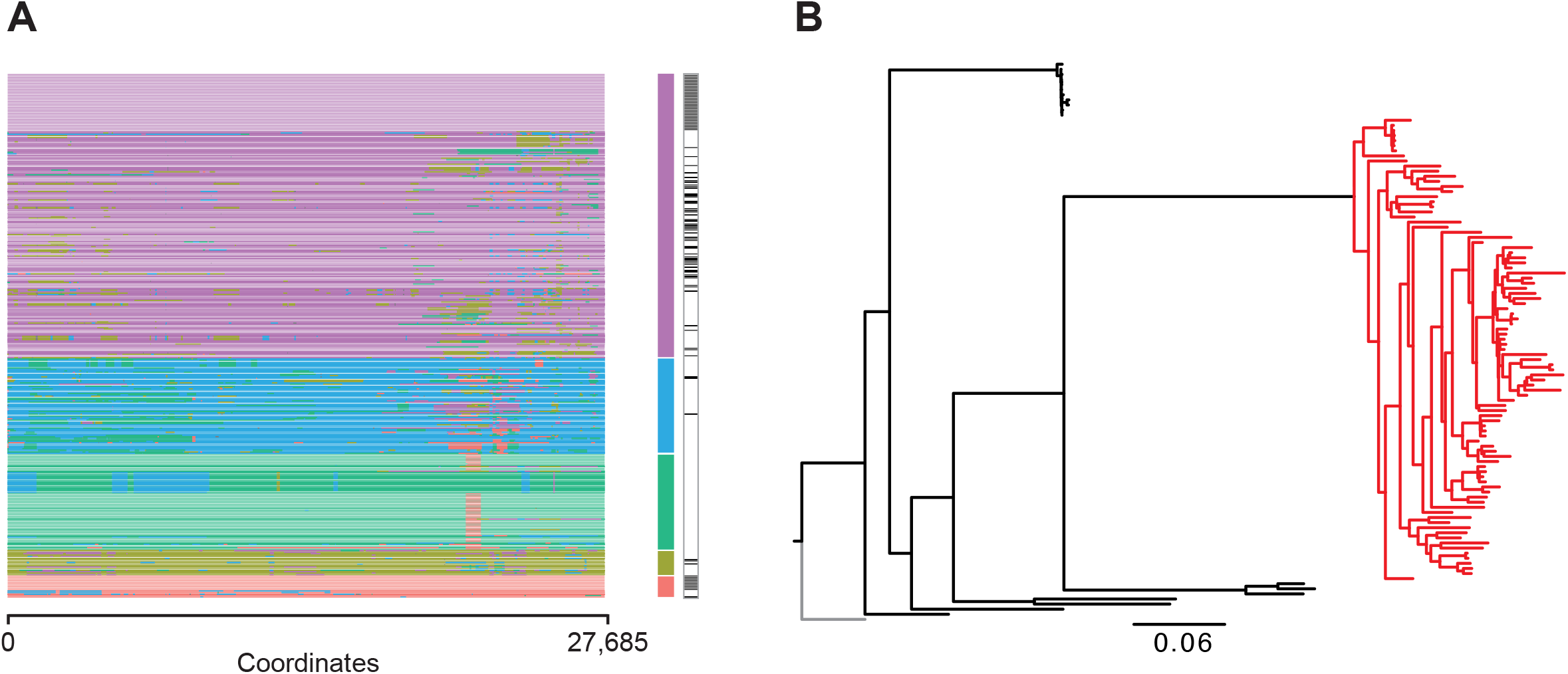
Representation of the steps we used to select strains for our metapopulation analysis through the example of the avian coronavirus. (A) Representation of the recent recombinations events happening within strains of the same metapopulation, inferred by fastGear. Rows correspond to viral strains sequences, the columns to positions in the alignment, and colors show the membership of each portion of sequences to lineage detected by fastGear. Sequence colored in black on the left side represent sequence selected to generate the phylogenetic tree. (B) Phylogenetic tree of the relationship among viral strains. The tree is rooted based on an outgroup (grey) as indicated on Supplementary Table 2.

**Supplementary Figure 4:**
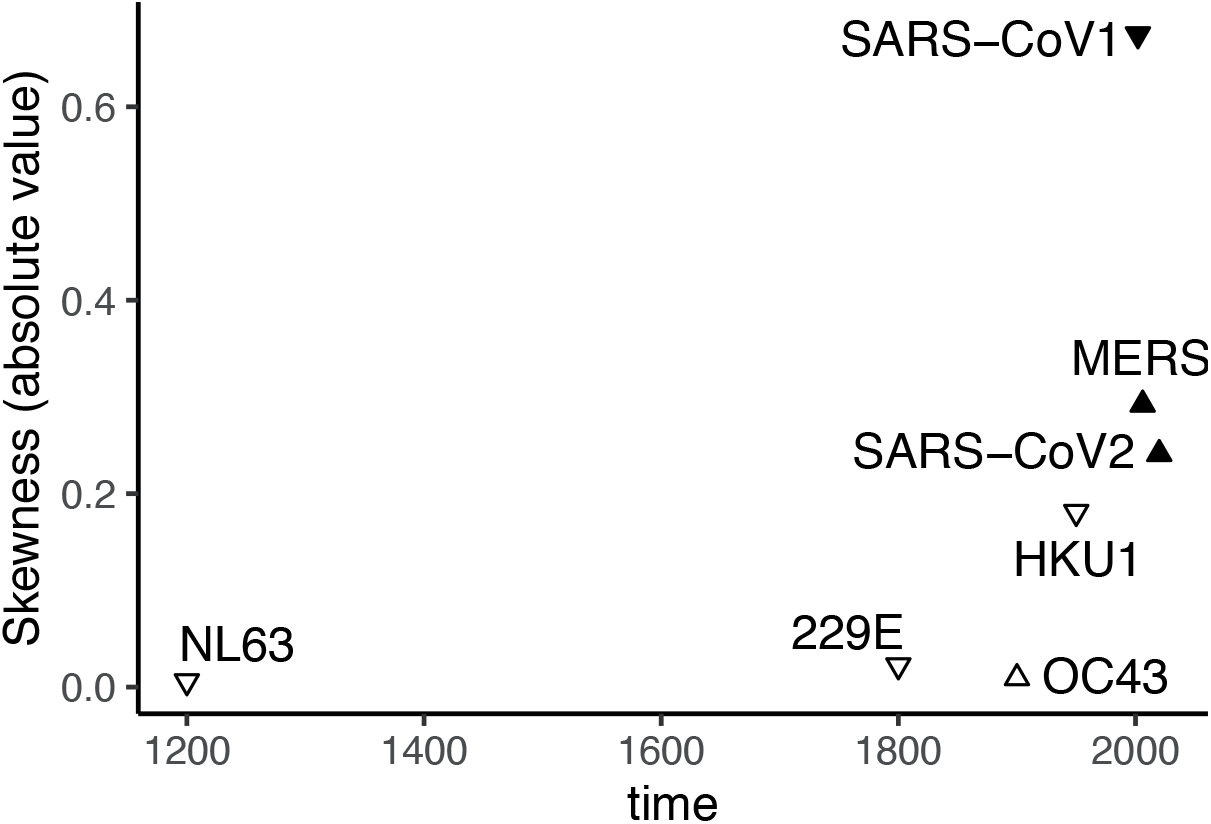
Scaterplot of the relationship between skweness of the folded SFS and coronavirus emergence time. The downward and upward triangle represented negative and positive skweness, respectively, and the filled triangle represented significantly different skweness. Principal component coordiantes

**Supplementary Figure 5:**
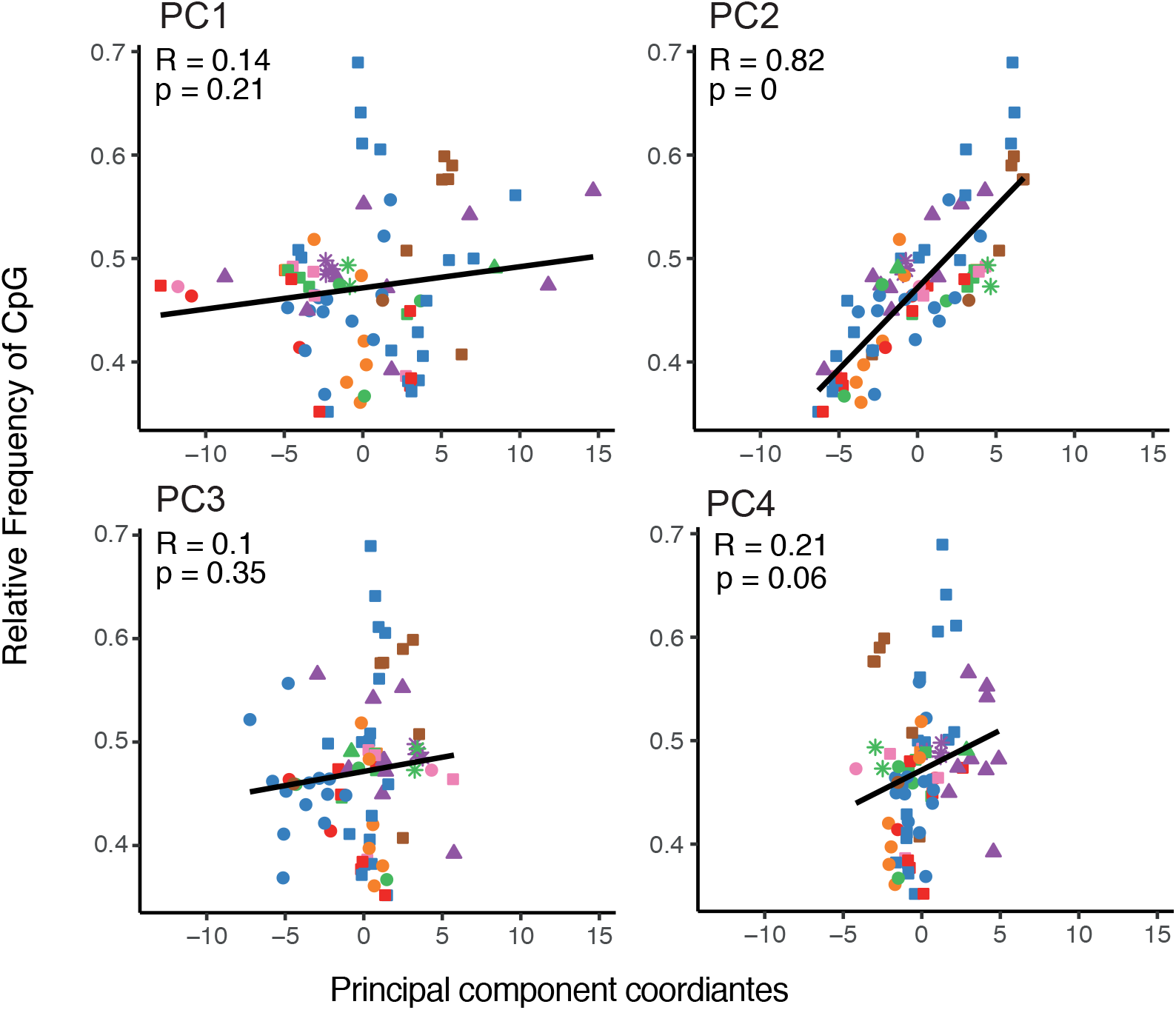
Regression between relative abundance of CpG dinucleotide and coordinate of the first 4 principal components. Principal component coordiantes

**Supplementary Figure 6:**
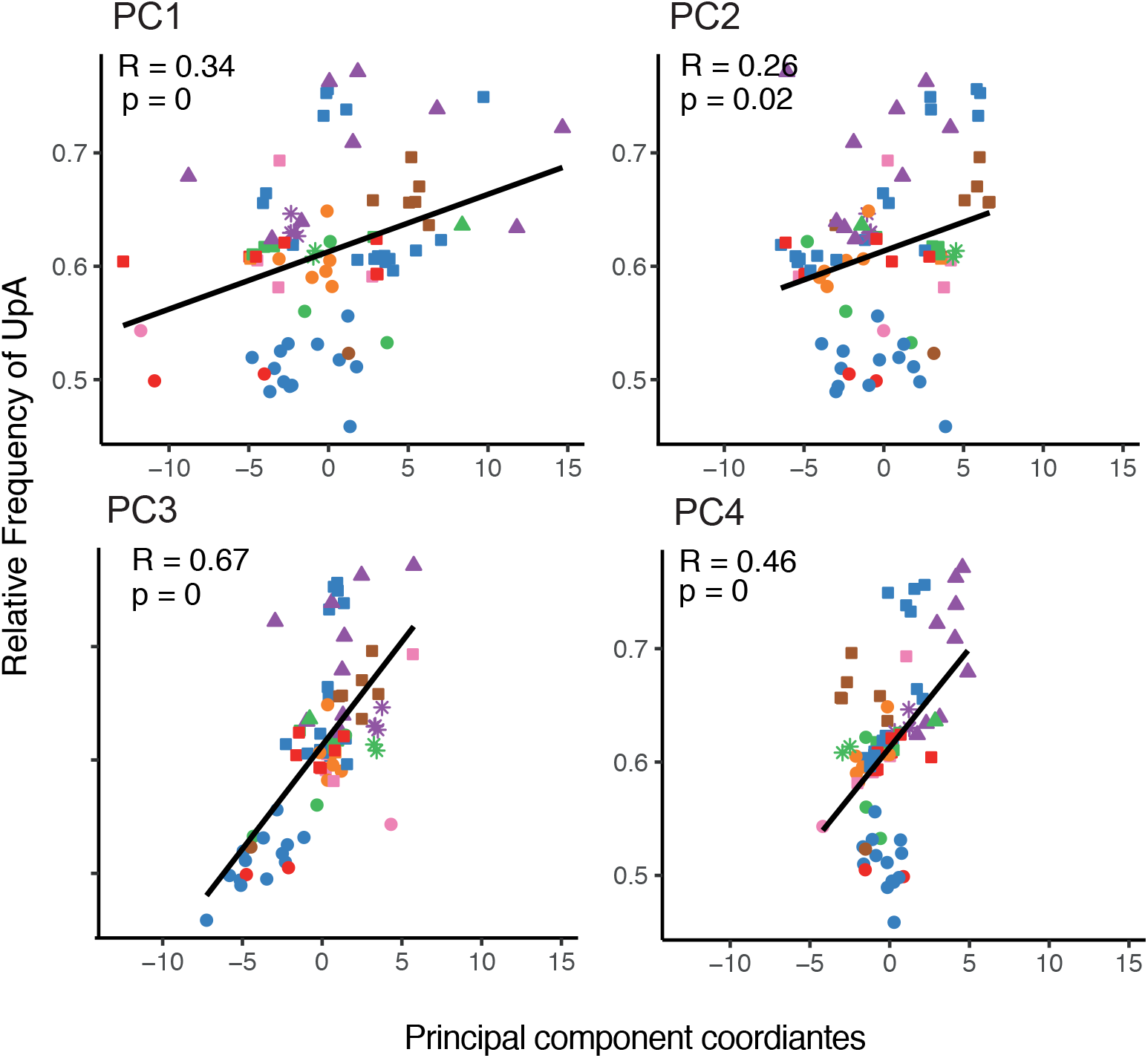
Regression between relative abundance of UpA dinucleotide and coordinate of the first 4 principal components.

**Supplementary Figure 7:**
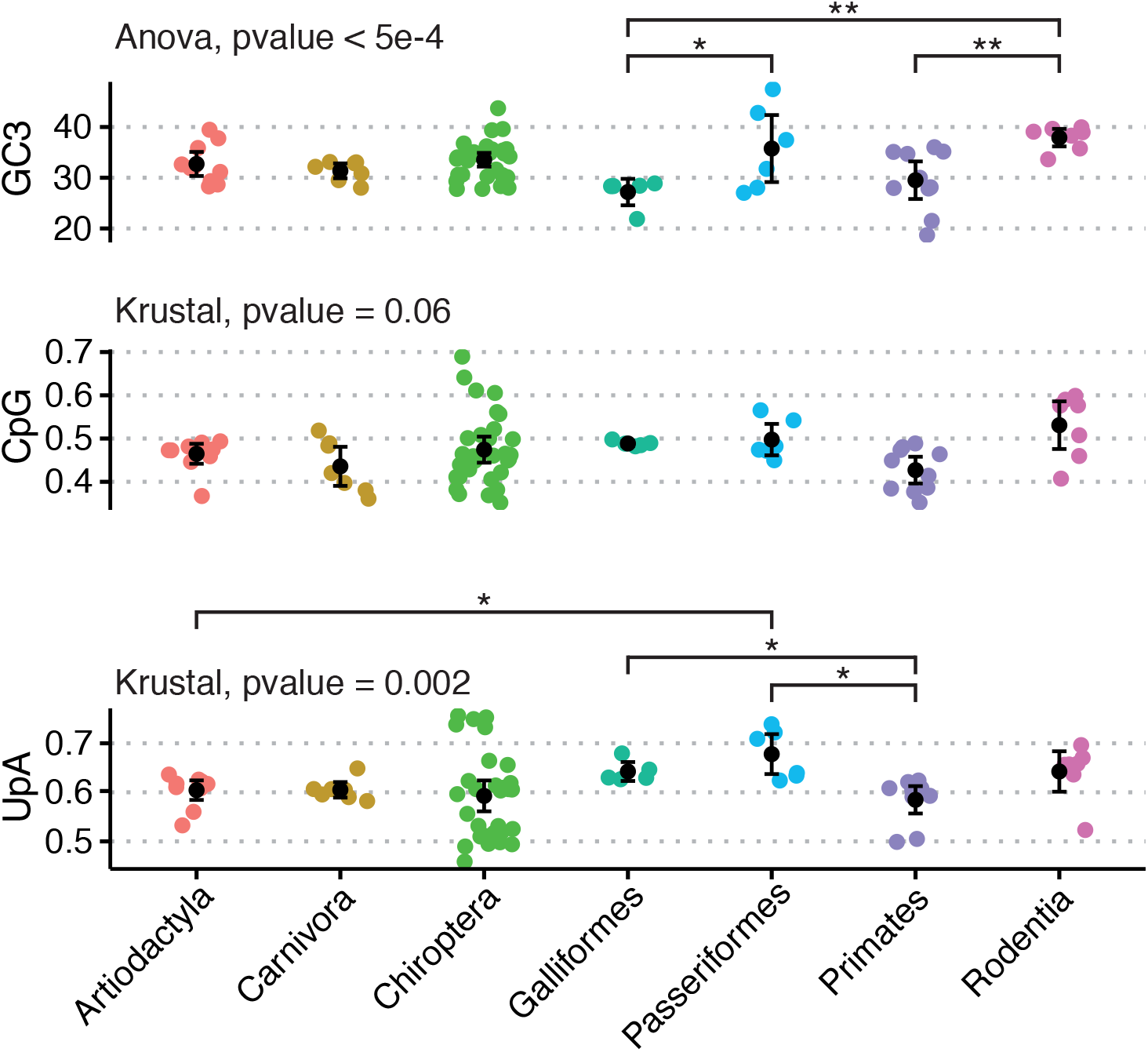
Comparison of the CG3 content, CpG, and UpA dinucleotide relative abundances across CoVs hosts. Variance analysis of normally distributed data was done using ANOVA follow by a post doc Tukey test showing further individual pairwise differences. Non-normal data were processed using Krustal-Wallis test follow by a rank Wilcoxon test with a Bonferroni correction (code for p-value: * <0.05, ** <0.01, *** <0.001).

**Supplementary Figure 8:**
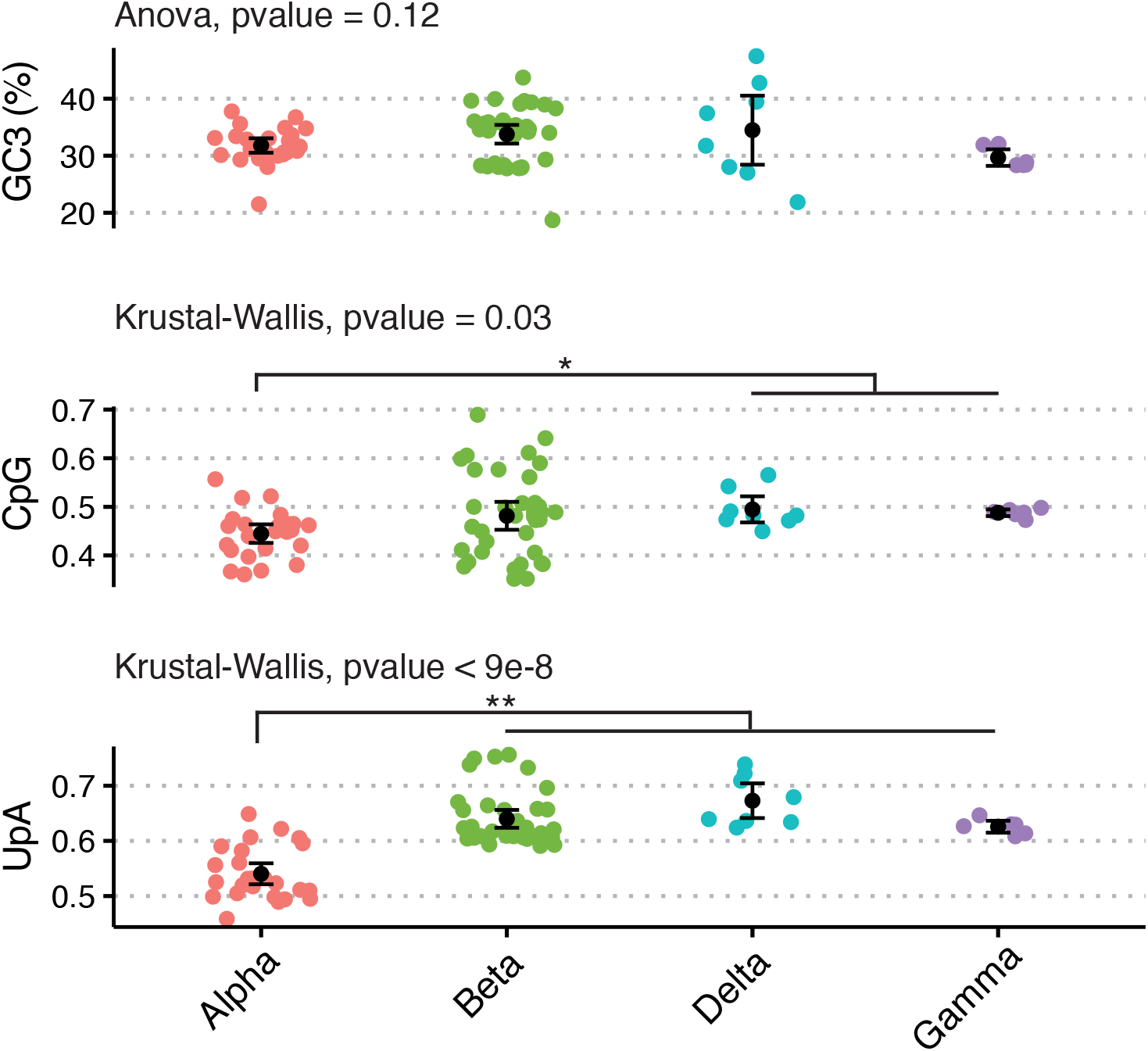
Comparison of the CG3 content, CpG, and UpA dinucleotide relative abundances across CoVs genera. Statistical significance has been treated similarly than above.

## Acknowledgements

The authors acknowledge the IRD itrop HPC (South Green Platform) at IRD Montpellier for providing HPC resources that have contributed to the research results reported within this paper. URL: https://bioinfo.ird.fr/-http://www.southgreen.fr. We warmly thank Sylvain Glémin for sharing with us R script function and helpful suggestions regarding our estimation of the mutational equilibrium.

## Funding

JD was supported by the Fondation pour la recherche Médicale (FRM, ARF20170938823) as well as by the Marie-Curie EU Horizon 2020 Marie-Skłodowska-Curie research and innovation program grant METHYVIREVOL (contract number 800489).

IGB was supported by the European Union Horizon 2020 research and innovation program under the grant agreement CODOVIREVOL (ERC-2014-CoG-647916).

## Author contributions

Conception: J.D. and I.G.B.; funding acquisition: J.D. and I.G.B.; method development and data analysis: J.D.; interpretation of the results: J.D. and I.G.B.; drafting of the manuscript: J.D. and I.G.B.

## Competing interests

The authors declare that they have no competing interests.

## Data Availability

We acknowledge the three different sources of genomes that we employed: the Virus-Host database (https://www.genome.jp/virushostdb/, version of March 19, 2020), the Virus Pathogen Resource database (VIRP, https://www.viprbrc.org/brc/home.spg?decorator=vipr, version of April 26, 2020), and the GISAID database (https://www.gisaid.org/, version of August 19, 2020).

## Notes

### Competing Interest Statement

The authors have declared no competing interest.

